# Interspecies differences in the transcriptome response of corals to acute heat stress

**DOI:** 10.1101/2024.08.16.607823

**Authors:** Jeric P. Da-Anoy, Niño Posadas, Cecilia Conaco

**Affiliations:** Marine Science Institute, University of the Philippines, Diliman, Quezon City, Philippines 1101; Department of Biology, Boston University, Boston, Massachusetts, USA 02215; Centre for Chromosome Biology, School of Biological and Chemical Sciences, University of Galway, Galway, Ireland

**Keywords:** thermal stress, adaptation, DNA damage, *Seriatopora caliendrum*

## Abstract

Rising sea surface temperatures threaten the survival of corals worldwide, with coral bleaching events becoming more commonplace. However, different coral species are known to exhibit variable levels of susceptibility to thermal stress events. To elucidate genetic mechanisms that may underlie these differences, we compared the gene complement of four coral species, *Favites colemani, Montipora digitata*, *Acropora digitifera,* and *Seriatopora caliendrum*, that were previously demonstrated to have differing responses to acute thermal stress. We found that more tolerant species, like *F. colemani* and *M. digitata,* possess a greater abundance of antioxidant protein families and chaperones. Under acute thermal stress conditions, only *S. caliendrum* showed a significant bleaching response, which was accompanied by activation of DNA damage response network and drastic upregulation of stress response genes (SRGs). This suggests that differences in SRG complement, as well as the mechanisms that control SRG expression response, contribute to the ability of corals to maintain stable physiological functions that is required to survive shifts in seawater temperature.

## Introduction

Coral communities worldwide are under threat due to the increasing sea surface temperature brought by changing global climate (Hughes et al. 2003; Hoegh-Guldberg et al. 2007; Carpenter et al. 2008;). Thermal anomalies in reefs has led to the increased frequency and severity of coral diseases and bleaching (Porter et al. 2001; Sutherland, Porter, and Torres 2004), consequently resulting to dramatic changes in reef community structure and general decline in coral cover (T. P. Hughes et al. 2003; Hoegh-Guldberg et al. 2007; De’ath et al. 2012). However, stress-tolerant corals may resist local perturbations and climate-change associated stressors, which may eventually repopulate reefs that have undergone mass bleaching events (van Woesik and Jordán-Garza 2011).

Corals exhibit differences in tolerance to thermal stress depending on species (Gibbin et al. 2015), morphology (Alvarez-Filip et al. 2011; McClanahan, Starger, and Baker 2015), or thermal history (Middlebrook, Hoegh-Guldberg, and Leggat 2008; Wall et al. 2018). For example, pocilloporids generally exhibit a low thermal threshold while an acroporid displays the highest overall tolerance among the Red Sea corals tested in a stress experiment (Evensen et al. 2022). Thermal tolerance of closely related species (e.g., genus *Acropora*) can, however, be divergent (Marshall and Baird 2000; Loya et al. 2001; Terry P. Hughes et al. 2017; Dalton et al. 2020), which may be attributed to the morphological diversity within a taxonomic group (Baird and Marshall 2002; Terry P. Hughes et al. 2017). Massive and encrusting colonies tend to be more resilient from bleaching compared to finely branched species (Loya et al. 2001). In the 1998 bleaching event on Ishigaki Island, massive *Porites* exhibited higher bleaching resilience than the branching morphotype (Kayanne et al. 2002). This variability in stress tolerance among corals is, in part, attributed to lineage-specific innovations in their molecular toolkit in responding to stress events. For example, comparative genomics of major scleractinian lineages showed that stress tolerance in corals is correlated to the number of genes encoding HSP20 proteins (Ying et al. 2018). A genomic survey of the starlet sea anemone, *Nematostella vectensis*, revealed that cnidarians have the components of stress response networks including genes engaged in responding to reactive oxygen species, toxic metals, osmotic shock, thermal stress, pathogen exposure, and wounding (Reitzel et al. 2008). Targeted comparison of cnidarian stress response gene (SRG) repertoire among diverse coral species within a reef may unveil other determinants of inter-species variability in coral stress tolerance.

Describing the acute and chronic stress response mechanisms in corals may also reveal short- and long-term acclimatization strategies, which may determine their adaptive capacity. For instance, the acute thermal stress response of coral larvae is accompanied by homeostatic functions (e.g., expression of heat shock proteins) while chronic stress response is associated with homeorhetic regulation (e.g., transcriptome-wide changes and expression shifts of translation machinery) (Meyer, Aglyamova, and Matz 2011a). This response was shown to be influenced by stress history, in which frequent exposure to stressful conditions can precondition coral populations which may enhance the thermal tolerance of even the susceptible groups (Bellantuono et al. 2012; Carilli, Donner, and Hartmann 2012). Branching *A. hyacinthus* population in a highly variable environment is thermally tolerant and exhibits high constitutive expression or frontloading of heat shock proteins and antioxidants, as well as genes involved in apoptosis, innate immunity, and cell adhesion (Barshis et al. 2013). This adaptation may represent genetically fixed acclimatization responses to recurring variable levels of temperature, pH, and oxygen (Craig, Birkeland, and Belliveau 2001; Smith et al. 2008) that may have persisted over several generations of the local coral population. Elucidating transcriptome-wide changes and expression patterns of SRGs in corals under stress may further uncover molecular mechanisms underlying differences in coral thermal tolerance.

Here, we asked how sympatric coral species in the Bolinao-Anda Reef Complex (BARC) in northwestern Philippines, a region that experiences thermal anomalies and steadily rising sea surface temperatures (Peñaflor et al. 2009; Yu 2012), would fare under thermal challenge. Four coral species, *Favites colemani, Montipora digitata, A. digitifera,* and *Seriatopora caliendrum*, were subjected to a common thermal regime. To examine changes in gene expression accompanying responses to thermal stress, we sequenced their transcriptomes and compared the expressed SRG complement. This genomic information provides a window into the differential susceptibilities of corals to elevated temperature and reveals the molecular mechanisms that may underlie these differences.

## Materials and Methods

### Coral collection and acclimatization

Three colonies each of *F. colemani, M. digitata, A. digitifera,* and *S. caliendrum* (Fig. 1A-D) were collected from depths of 2-9 m within the Bolinao-Anda Reef Complex in November 2016 (Table S1). Sea surface temperatures within the reef range from 25-32 °C with an annual mean temperature of 28.89±0.90 °C based on monitoring data from the Bolinao Marine Laboratory. Samples were collected with the permission of the Philippines Department of Agriculture Bureau of Fisheries and Aquatic Resources (DA-BFAR Gratuitous Permit No. 0102-15). Colonies at least 10-15 m apart were collected to minimize genotypic similarity, although sample genotypes were not evaluated. Corals were fragmented into 2.5-5.0 cm long nubbins and pre-conditioned for two weeks in outdoor tanks with running seawater maintained at 28±1 °C and illumination under low photosynthetic photon flux density (∼80-90 µmol m^-2^ s^-1^) on a 12:12 light-dark cycle. Fragments were tagged to keep track of their colony of origin. Healed fragments were then allowed to acclimatize for two weeks in indoor experimental tanks with running seawater maintained at 28±1 °C and illumination of ∼80 µmol m^-2^ s^-1^ on a 12:12 light-dark cycle.

**Fig 1.**
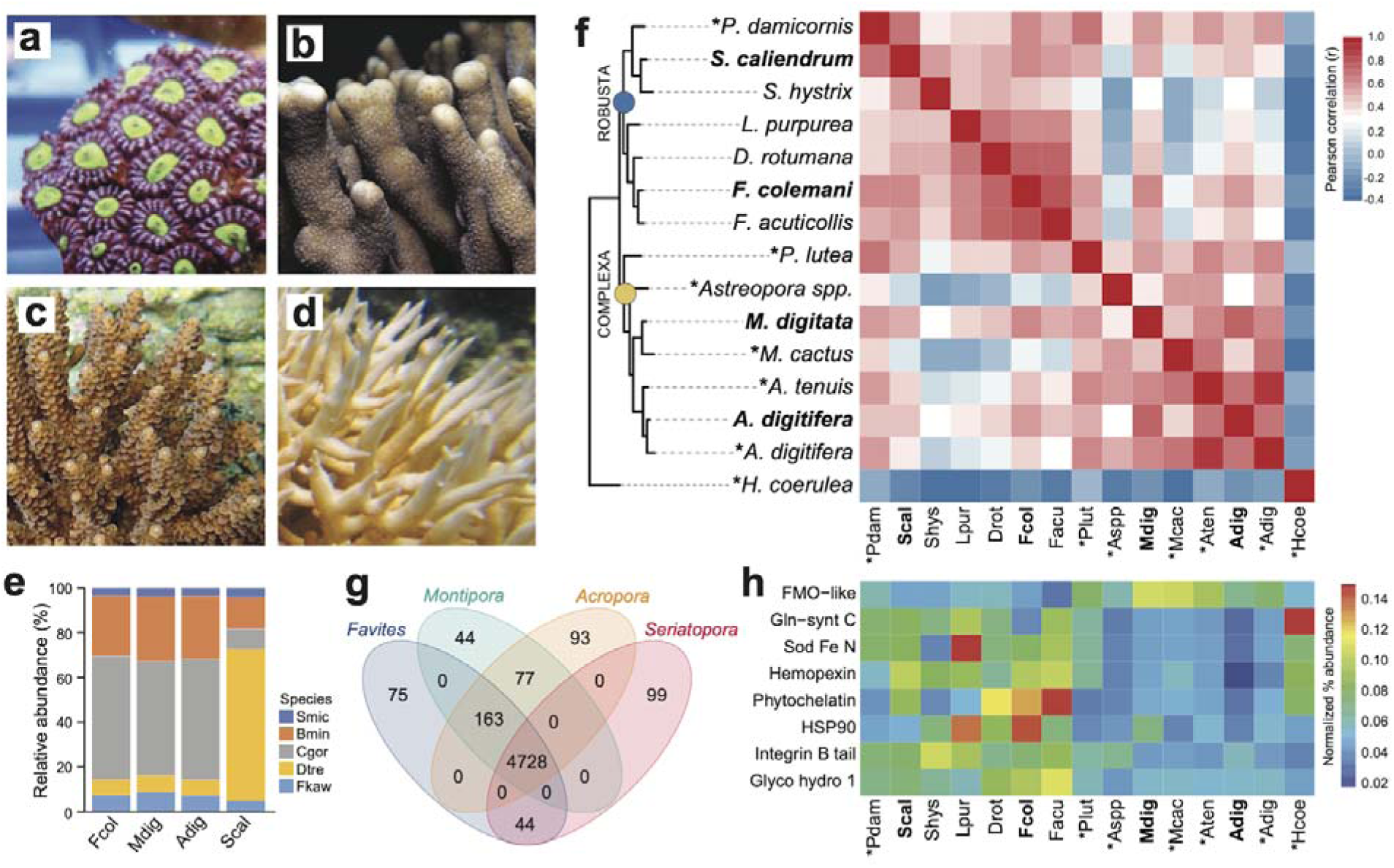
Comparison of coral *de novo* transcriptomes. Representative photographs of (A) *Favites colemani*, (B) *Montipora digitata*, (C) *Acropora digitifera*, and (D) *Seriatopora caliendrum.* (E) Relative abundance of symbiont transcripts with best hits to genomes of other Symbiodiniaceae. Smic, *Symbiodinium microadriaticum*; Bmin, *Breviolum minutum*; Cgor, *Cladocopium goreaui*; Dtre, *Durusdinium trenchii*; Fkaw, *Fugacium kawagutii*. (F) Orthologous groups in coral host transcriptomes from this study (in bold font) and in genomes (asterisks) or transcriptomes of other coral species from the Robusta (blue circle) and Complexa (yellow circle) superfamilies. Heatmap colors represent Pearson’s correlation coefficient for orthogroups between species pairs. (G) Number of unique and common PFAM domains identified in the coral host transcriptomes. (H) Enriched PFAMs across different coral species.

### Thermal stress experiments

Thermal stress experiments were conducted in 40 L tanks supplied with constantly aerated, 10 μm-filtered flow-through seawater, as described in (Da-Anoy, Cabaitan, and Conaco 2019)). Briefly, after two weeks of acclimation, fragments were transferred directly to experimental tanks set at 28 °C (control) or 32 °C (heated). Two independent replicate tanks were used for each temperature treatment. Each treatment tank contained at least 15 coral fragments (with 5-6 fragments from each of the three colonies of the four coral species used in this study).

Seawater was pumped into treatment tanks from chilled reservoirs at a rate of 5-8 L/hr. The temperature in each tank was adjusted to the target setting using submersible thermostat heaters (EHEIM GmbH & Co. KG, Baden Wurttemberg, Germany) with a water pump (600 L/hr) to augment circulation. All setups were illuminated under low photosynthetic photon flux density (∼80 µmol m^-2^ s^-1^) on a 12:12 light-dark cycle to avoid light stress. Temperature and light intensity were monitored using submersible loggers (HOBO pendant, Onset Computer Corp., Bourne, MA, USA). Coral fragments were collected after 4 and 24 hr exposure to treatment conditions. Samples were flash-frozen in liquid nitrogen for transport and then stored at -80 °C before processing.

### RNA extraction and sequencing

Total RNA was extracted using TRIzol Reagent (Invitrogen, Waltham, MA, USA) following the manufacturer’s protocol. Contaminating DNA was removed using the DNA-free kit (Invitrogen). RNA integrity of samples were determined by electrophoresis on a native agarose gel with denaturing loading dye. Quantitation of RNA was done using the BioSpec Nanodrop spectrophotometer (Shimadzu, Kyoto, Japan). Samples with an RNA integrity number (RIN) below 7.8 were excluded from sequencing. Three biological replicates representing RNA from control and heated treatments collected at 4 and 24 hr durations (12 samples per species) were sent to BGI Genomics, Hong Kong, or Macrogen, South Korea, for library preparation and sequencing. mRNA enrichment and preparation of barcoded cDNA libraries were done using the TruSeq RNA Sample Prep Kit (Illumina, Inc., San Diego, CA, USA). *F. colemani* and *M. digitata* were sequenced on the HiSeq 4000 platform (Illumina, Inc.) at Macrogen, while *A. digitifera* and *S. caliendrum* libraries were sequenced on the HiSeq 2500 platform (Illumina, Inc.) at BGI Genomics, to generate 100 bp paired-end reads.

### Preprocessing of reads, transcriptome assembly, and annotation

Raw sequence read quality was assessed using FastQC 0.10.1 (Andrews 2010) and trimmed through Trimmomatic 0.32 (Bolger, Lohse, and Usadel 2014). Poor-quality bases (quality score < 3) at leading and trailing bases, as well as the first 15 bases of the reads were discarded. Reads less than 36 bases long and with an average quality per base (4-base sliding window) < 30 were trimmed. *De novo* transcriptome assembly was carried out through Trinity (Grabherr et al. 2011) using eight libraries for each species. The assembled reference transcriptomes were subjected to CD-HIT-EST (Fu et al. 2012) to cluster genes at 90% identity. To further reduce assembly redundancy, transcript abundance estimation was performed by mapping trimmed reads back to the reference transcriptomes using RNASeq by Expectation-Maximization (RSEM) (B. Li and Dewey 2011) and the Bowtie alignment method (Langmead et al. 2009). Transcript isoforms with zero IsoPct were removed. Isoforms with the highest combined IsoPct or longest length were retained for each transcript. Protein-coding regions for each transcript were predicted using TransDecoder v.2.0.1 (https://github.com/TransDecoder/transdecoder) package in Trinity. Only protein-coding transcripts and their longest predicted peptide sequences were retained for subsequent analyses.

To segregate host and symbiont sequences, Psytrans (https://github.com/sylvainforet/psytrans) was implemented using curated peptide sequence databases of corals (i.e., *Montipora capitata* (Shumaker et al. 2019), *A. digitifera* (Shinzato et al. 2011), *Goniastrea aspera, Galaxea fascicularis* (Ying et al. 2018), *A. tenuis,* and *Porites lutea* (ReFuGe 2020 Consortium 2015) and Symbiodiniaceae representatives (i.e., *Fugacium kawagutii, Cladocopium goreaui (T. Li et al. 2020)*, *Breviolum minutum*, *Cladocopium* sp., *Symbiodinium* sp. (Shoguchi et al. 2013), and *Cladocopium* C15 (Robbins et al. 2019), respectively. To further remove potential contaminating sequences from other epibionts, predicted peptides were aligned by Blastp (e-value ≤ 10^−5^) against the Genbank non-redundant (nr) sequence database. Only sequences with a best match to Cnidaria (for host) or Dinophyceae (for symbiont) were retained in the final non-redundant reference assemblies. Assembly completeness was assessed through Benchmarking Universal Single-Copy Orthologs (BUSCO) (Simão et al. 2015) using the Eukaryota, Metazoa, and Alveolata ortholog databases.

The final non-redundant reference transcriptomes were annotated following the Trinotate pipeline (https://trinotate.github.io/). Homolog search was performed by aligning nucleotide and predicted peptide sequences against the UniProt/Swiss-Prot (UniProt 2019), Genbank RefSeq (O’Leary et al. 2016), and nr databases (Pruitt, Tatusova, and Maglott 2005) through Blastx and Blastp (e-value ≤ 10^−5^). Protein domains were identified through HMMER v.3.1b2 (http://hmmer.org) using the Pfam-A database (v31.042). The top Blast hits and identified protein domains for each gene were used as inputs into Trinotate to predict gene ontology (GO) annotations and to generate a comprehensive assembly annotation report.

### Ortholog analysis, gene content comparison, and symbiont sequence similarity

Orthologous gene families in the host transcriptomes of *F. colemani, M. digitata, A. digitifera,* and *S. caliendrum* and in the genomes or transcriptomes of other coral species were identified using OrthoFinder (Emms and Kelly 2019). A total of 10 other coral species representing superfamily Robusta (*P. damicornis*, *S. hystrix*, *Leptastrea purpurea*, *Dipsastrea rotumana*, and *F. acuticollis*) and superfamily Complexa (*Porites lutea*, *Astreopora* spp., *M. cactus*, and *A. tenuis*) were included in the analysis, with an octocoral (*Heliopora coerulea*) as outgroup (Table S2). The classification of these species into the Robusta and Complexa superfamilies was based on the works of (Zhang et al. 2019; Okubo 2016) and (Ying et al. 2018). Enriched orthologous genes among taxonomic groups were identified using KinFin (Laetsch and Blaxter 2017). Predicted peptide sequences of representative coral species were annotated against the Pfam 32.0 (Finn et al. 2014) database to annotate expanded orthogroups, as well as to identify lineage-restricted protein domains.

Symbiont transcriptomes from *F. colemani, M. digitata, A. digitifera,* and *S. caliendrum* were aligned using Blastp at an e-value cutoff of 1 × 10^−5^ against genomes of other Symbiodiniaceae representatives, including *S. microadriaticum*, *B. minutum*, *C. goreaui*, *Durusdinium trenchii*, and *F. kawagutii* (Table S2). The affiliation of each transcript was assigned based on its top Blastp hits (highest % identity and lowest e-value).

### Identification of stress response genes and transcription regulators

Cnidarian stress response genes (SRGs) (Reitzel et al. 2008), as well as gene regulatory elements, including transcription factors (Bahrami, Ehsani, and Drabløs 2015) and epigenetic modifiers (Lee and Workman 2007; Kooistra and Helin 2012; Seto and Yoshida 2014; de Mendoza et al. 2019), were identified in the host transcriptomes of *F. colemani, M. digitata, A. digitifera,* and *S. caliendrum* based on their characteristic domains and top Blastp hit (e-value < 1x10^-5^) against the UniProtKB/Swiss-Prot database.

The abundance of SRGs was computed relative to the total number of predicted peptides in each species. SRGs that distinguish between coral taxonomic groups were identified using the Multiple Variable Associations with Linear Models (MaAsLin 2) (Mallick et al. 2021) package implemented in R.

### Differential gene expression analysis

Trimmed reads were mapped back to the concatenated host and symbiont reference transcriptomes to estimate transcript abundance using RSEM (B. Li and Dewey 2011) and the Bowtie alignment method (Langmead et al. 2009). Expected counts were converted to counts per million (CPM) and only genes with > 2 CPM in at least two libraries (*F. colemani* = 43 336, *M. digitata* = 43 320, *A. digitifera* = 49 343, *S. caliendrum* = 44 171) were included in differential gene expression analysis. Time-matched pairwise comparisons between control and heated samples were conducted using the edgeR (Robinson, McCarthy, and Smyth 2010) package in R. Transcripts with a log_2_ fold change (log_2_FC) ≥ |4| and a false discovery rate (FDR)-adjusted *p*-value < 0.05 were considered differentially expressed. GO enrichment analysis for differentially expressed genes (DEGs) was performed using the topGO package (Alexa and Rahnenführer 2009) in R. Only GO terms with *p*-value < 0.01 were considered significantly enriched. Enriched terms were summarized through REVIGO (Supek et al. 2011) at 0.5 similarity cut-off.

Predicted peptides of *S. caliendrum* were searched against the human proteome v.11.5 from the STRING v.11 database (Mering et al. 2003) with an e-value cut-off of 1x10^-5^. Blastp top hits for DEGs in either 4- or 24-hr comparisons were used as input in pathway enrichment analysis. Protein-protein interactions of genes involved in enriched pathways (score > 0.400 and FDR < 0.01) were retrieved from the STRING v.11 database (Mering et al. 2003). Interaction networks were visualized using Cytoscape v.3.7.2 (Shannon et al. 2003). Relative expression of *S. caliendrum* gene homologs in heated samples relative to the controls was computed as the average sum of transcripts per million (TPM).

To assess the effect of treatments, raw counts of host- and symbiont-derived transcriptomes were rlog-transformed and used as input for principal component analysis (PCA) with `plotPCA` (DESeq2 package), followed by PERMANOVA using the `adonis2` function from the vegan package (Oksanen et al. 2022). Gene expression plasticity in response to treatments was calculated following (Bove et al. 2023). Plasticity was calculated as the distance between an individual’s transcriptome profile and the mean of all samples in the 4 hr control group. Differences in plasticity between treatments were tested using an ANOVA followed by Tukey’s HSD post-hoc tests.

### Visualization

All visualizations were done using the ggplot2 package (Wickham 2016) in R. Phylogenetic trees were edited in iTOL (Letunic and Bork 2019).

## Results

### *De novo* assembly of four scleractinian coral transcriptomes

High-throughput sequencing of *F. colemani*, *M. digitata*, *A. digitifera*, and *S. caliendrum* (Fig. 1A-D) transcriptomes generated 368.74 to 688.08 M raw reads per species (Table S3). Quality-filtered reads were assembled *de novo* (Table S1). Resulting assemblies were subjected to sequence similarity clustering, isoform selection, and removal of non protein-coding and non-Cnidaria or non-Dinophyceae sequences to generate non-redundant reference transcriptomes for each species (*F. colemani*: n = 52 832, Ex90N50 = 2 123; *M. digitata*: n = 51 324, Ex90N50 = 2 045; *A. digitifera*: n = 65 543, Ex90N50 = 2 379; *S. caliendrum*: n = 54 146, Ex90N50 = 2 346) (Table 1, Table S1). Assemblies contained a greater proportion of symbiont (48.25 - 55.93%, GC content = 54.94 - 56.04%) compared to host transcripts (44.07 - 51.75%, GC content = 41.78 - 42.82%) (Table 1, Table S1). BUSCO analysis revealed that the host assemblies were about 90.90 - 93.80% complete for metazoan core genes and the symbiont assemblies were 66.70 - 70.80% complete for alveolate core genes (Table 1, Table S4). Around 67.08 - 73.41% of host transcripts were annotated by at least one of the following databases: UniProtKB/Swiss-Prot, PFAM, or GO. Symbiont assemblies had a relatively lower annotation rate of 54.92 - 57.90% (Table S5).

**Table 1.** Assembly statistics of *de novo* transcriptomes of four scleractinian corals.

|  | *Favites colemani* | *Montipora digitata* | *Acropora digitifera* | *Seriatopora caliendrum* |
| --- | --- | --- | --- | --- |
| Superfamily | Robusta | Complexa | Complexa | Robusta |
| Family | Merulinidae | Acroporidae | Acroporidae | Pocilloporidae |
| Total transcripts | 52 832 | 51 324 | 65 543 | 54 146 |
| Total transcripts ExN50 | 2 123 | 2 045 | 2 379 | 2 346 |
| **Host bin** |  |  |  |  |
| Number of Transcripts | 25 681 | 22 616 | 33 918 | 24 763 |
| G+C content (%) | 42.82 | 42.51 | 42.03 | 41.78 |
| % completeness (BUSCO Metazoa db) | 92.00% | 93.00% | 93.80% | 90.90% |
| **Symbiont bin** |  |  |  |  |
| Number of Transcripts | 27 151 | 28 708 | 31 625 | 29 383 |
| G+C content (%) | 55.03 | 55.24 | 54.94 | 56.04 |
| % completeness (BUSCO Alveolata db) | 70.80% | 69.00% | 67.30% | 66.70% |

### Symbiont transcript affiliations

Symbiont assemblies were aligned by Blastp to genomes of other Symbiodiniaceae to determine the possible identities of microalgal symbionts associated with the four coral hosts. About 51.07-55.26% of symbiont transcripts in *F. colemani*, *M. digitata*, and *A. digitifera* had a best hit with sequences from *C. goreaui* and 6.84-7.45% to *D. trenchii* (Fig. 1E, Table S6). In contrast, only 9.10% of symbiont transcripts in *S. caliendrum* had a best hit with *C. goreaui*, while 67.69% showed highest similarity with sequences from *D. trenchii*.

### Comparison of coral host assemblies

Ortholog analysis was conducted to assess gene conservation across coral species and to explore possible gene expansion events. The gene repertoire of our coral host assemblies were comparable to other coral transcriptomes or genomes, indicating that we were able to capture most of the scleractinian core genes. Correlation of orthologous groups that were identified across species recapitulated phylogenetic groupings, with *S. caliendrum* and *F. colemani* clustering with other members or suborder Vacatina (Robusta superfamily) and *M. digitata* and *A. digitifera* clustering with other members of suborder Refertina (Complexa superfamily) (Fig. 1F, Table S7). The majority of transcripts from our assemblies are conserved in Anthozoa (4 296 orthogroups) and scleractinian representatives (740 orthogroups) (Table S8). Forty three orthogroups are represented in Robusta and 17 in Complexa. Family-specific orthologous genes were also identified among representatives of pocilloporids (n = 57), merulinids (n = 44), and acroporids (n = 47).

Ortholog-based and taxon-aware analyses using KinFin (Laetsch and Blaxter 2017) revealed genes that had undergone lineage-specific expansions (Table S9). An ion transport protein (OG224) and secretin GPCR (OG190) are relatively enriched in members of family Pocilloporidae, while HSP70 (OG128), NHL repeat (OG141 and OG236), B-box zinc finger (OG134), and carboxylesterases (OG233) are expanded in merulinid species. A total of 30 orthogroups are enriched among acroporids, including rhodopsin GPCR (OG254), ubiquitin transferase (OG178), DNA-binding THAP (OG13), reverse transcriptases (OG38, OG0, and OG205), and helicases (OG145 and OG179), as well as a diversity of integrase (OG10), transposases (OG229, OG71, OG49, OG78, OG73, and OG123), and endonucleases (OG44, OG230, OG222, OG110, and OG281). Immune-related orthogroups, such as NOD-like receptors (OG29), immunoglobulin (OG76 and OG156), and TRAF-type zinc finger (OG274), were also enriched among members of genus *Acropora*.

### Gene complement and SRG repertoire

To evaluate the functional gene complement represented in each species, we compared protein domains identified in the translated transcriptomes of *Favites*, *Montipora*, *Acropora*, and *Seriatopora*. This revealed 4 728 domains conserved in all four species. 163 protein domains were shared between *Favites*, *Montipora* and *Acropora* (n = 163), including an ABC transporter family (PF06541), flavoprotein (PF02441), DNA damage repair protein (PF08599), PAC3 (PF10178), Pcc1 (PF09341) and heat shock transcription factor (PF06546)) (Fig. 1G, Table S10). Domains restricted to *Seriatopora* and *Favites* (n = 44) included players involved in fungal-like histidine biosynthetic pathway (i.e., HisG (PF01634), HisG_C (PF08029), Histidinol_dh (PF00815), IGPD (PF00475), PRA-CH (PF01502), and PRA-PH (PF01503)), as well as ectoine synthase (PF06339), selenoprotein P (PF04592), an ER-bound oxygenase (PF09995), FOXO-TAD (PF16676), and transmembrane protein families (PF3616, PF07114, PF16070, and PF15475) (Table S10). On the other hand, protein domains specific to *Montipora* and *Acropora* (n = 77) included cell cycle regulatory protein Spy1 (PF11357), serpentine type GPCR (PF10318), DNA binding proteins (HTH_32 (PF13565), MBF2 (PF15868), TBPIP (PF07106), and Tfb5 (PF06331), cutA1 divalent ion tolerance protein (PF03091), membrane transport protein (PF03547), HSF binding factor (PF06825), stress-tolerance associated domain CYSTM (PF12734), Sep15/SelM redox domain (PF08806), zinc fingers (PF02892, and PF11781), and protein domains linked to DNA damage response and repair (UPF0544 (PF15749) and TAN (PF11640)). A total of 99 domains were restricted to *Seriatopora (*e.g., E3 ubiquitin ligases (PF12185 and PF10302), ROS modulator Romo1 (PF10247), odorant receptors (PF02949), and NF-κB modulator NEMO (PF11577)), 75 to *Favites* (e.g., C2H2 type zinc finger (PF13909), oxidoreductase (PF07914), and serpentine chemoreceptor (PF07914)), 44 to *Montipora* (e.g., serpentine chemoreceptor (PF10292)), and 93 to *Acropora* (e.g., cell wall stress sensor (PF04478) and peroxidase (PF00141)).

To gain insights into the stress response gene (SRG) repertoire of each coral species, we identified known cnidarian SRGs involved in chemical, pathogen, and wounding stress (Rietzel et al., 2008). Most SRG families were represented in each species (Table S11). Proteins engaged in chemical stress response (i.e., HSP90, phytochelatin, and Fe/Mn SOD) and pathogen defense (i.e., glycosyl hydrolase) were enriched in members of superfamily Robusta, particular in the merulinids, while proteins with flavin-binding monooxygenase and caspase recruitment domains were relatively more abundant in members of superfamily Complexa (Fig. 1H, Table S12-13).

### Transcriptome response to elevated temperature

To determine how the different coral species respond to the same thermal stress regime, we subjected coral fragments to acute elevated temperature (32°C vs 28°C) for 4 and 24 hr. Principal component analysis showed shifts in global gene expression profiles between heated and control samples for host-derived transcripts in *A. digitifera*, *F. colemani*, and *S. caliendrum* (Fig. 2A-D), as well as for symbiont-derived transcripts in *M. digitata* and *S. caliendrum* (Fig. S1A-D). This shift was quantified using gene expression plasticity analysis, which showed a significant difference in treatments only in *S. caliendrum* (*p*-value ≤ 0.05), indicating significantly higher plasticity in heated treatments, particularly at 24 hr exposure (Fig. 2A-D).

**Fig 2.**
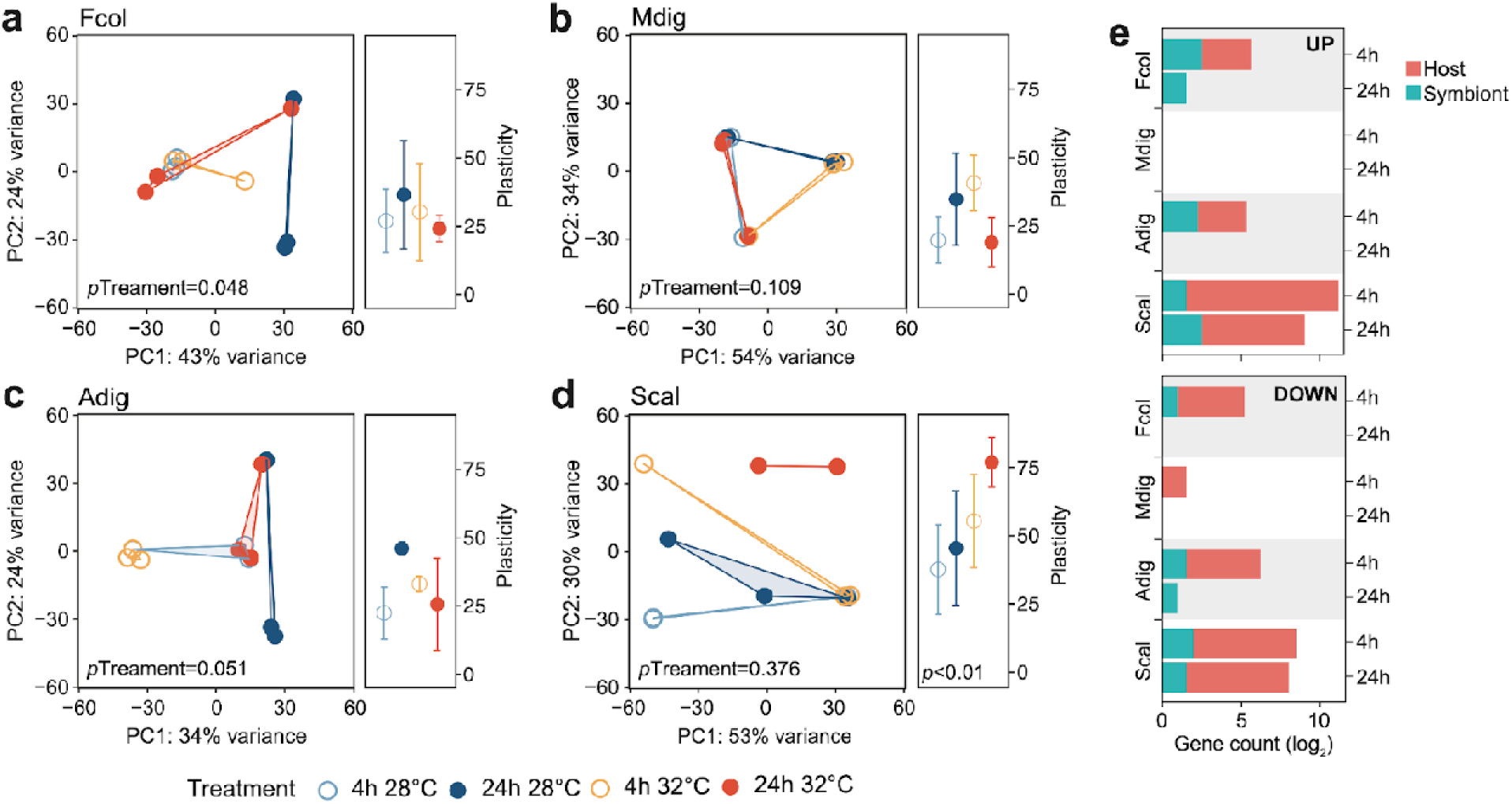
Transcriptome dynamics under thermal stress. Principal component analysis (PCA) and gene expression plasticity plots of host-derived transcriptomes for (A) *F. colemani*, (B) *M. digitata*, (C) *A. digitifera*, and (D) *S. caliendrum* in different treatments. The x- and y-axes represent the % variance explained by the first two principal components. (E) Differentially expressed (log_2_FC ≥ |4|, FDR-adjusted *p*-value < 0.05) host and symbiont genes (upregulated, top; downregulated, bottom) after 4 hr and 24 hr exposure.

Differential expression analysis support this finding, with fewer genes exhibiting a significant change in expression (log_2_FC ≥ |4|, FDR ≤ 0.05) in *F. colemani* (94 at 4 hr, 2 at 24 hr), *M. digitata* (2 at 4 hr, 0 at 24 hr), and *A. digitifera* (124 at 4 hr, 1 at 24 hr) subjected to elevated temperature (Fig. 2E, Table S14). In contrast, *S. caliendrum* had 2 865 differentially expressed transcripts at 4 hr (2 495 upregulated, 370 downregulated), and 823 transcripts differentially expressed at 24 hr (559 upregulated, 263 downregulated). Relative to time-matched controls, more transcripts were differentially expressed at 4 hr compared to the 24 hr timepoint in all species. Majority of differentially expressed transcripts originated from the coral host.

Host genes upregulated in *F. colemani* subjected to acute thermal stress included antioxidants (*MOXD1* and *PXDNL*), solute carrier proteins (*S6A13*, *SO4A1*, and *COPT2*), and ABC transporters (2 *MRP4*), as well as ECM components (*HMCN2*, 2 *SNED1*, *SUSD2*, and *MYO1*), whereas heat shock proteins (*HSP7C* and *HSP16*) and ubiquitination-related genes (*TRI50* and *UBC12*) were downregulated (Table S15). In *A. digitifera*, oxidoreductase (*QORL1*) and ECM-associated genes (*ITAD* and *MDGA2*) were upregulated, whereas *HSP7A*, E3 ubiquitin protein ligases (*R113A* and *TRAF6*), and translation initiation factors (2 *IF4G1*) were downregulated (Table S16).

### *Seriatopora caliendrum* thermal stress response

In contrast to the other corals that showed a minimal transcriptional response to the experimental treatment, *S. caliendrum* exhibited a significant shift in expression of genes involved in diverse biological processes, including basic cellular functions (e.g., growth, cell cycle, cell communication, cell adhesion, and DNA replication), metabolism (e.g., glyoxylate cycle, carbohydrate metabolism, nitrogen metabolism, DNA biosynthesis, cholesterol catabolism, and xenobiotic metabolism), gene expression control (e.g., microRNA-mediated gene silencing, DNA methylation, and post-translational modification), immune response (e.g., endocytosis, bacterial agglutination, and interferon-beta production), and stress response pathways (e.g., mismatch repair, heat acclimation, response to decreased oxygen levels, oxygen radical, and pH) (Table S17, Table S18). These broadscale transcriptional changes were accompanied by dynamic expression of gene regulatory elements such as epigenetic modifiers (DNA methyltransferases (*DNMT1* and *DNM3A*), thymine DNA glycosylase (*UNG* and *TDG*), histone deacetylase (*HDAC6*), methyltransferases (*PRD14*, *PRDM6*, and *SMYD3*), and demethylases (*KDM8*, *JMJD4*, and *KDM1B*)), along with diverse transcription factors (*ARID*, *bZIP*, *E2F*, *Ets*, *Forkhead*, *GATA*, *H2TH*, *bHLH*, *HMG box*, *Homeobox*, *Myb*, *Pou*, *T-box*, *THAP*, and zinc finger factors) (Fig. 3A, Table S19). The dynamic expression of these regulators signal active involvement of transcriptional control in the thermal stress response of *S. caliendrum*. Most of these regulators exhibited notable change in expression at 4 hr and returned to their basal levels after 24 hr (Fig. 3A), coinciding with observed transcriptome-wide dynamics (Fig. 2A-E).

**Fig 3.**
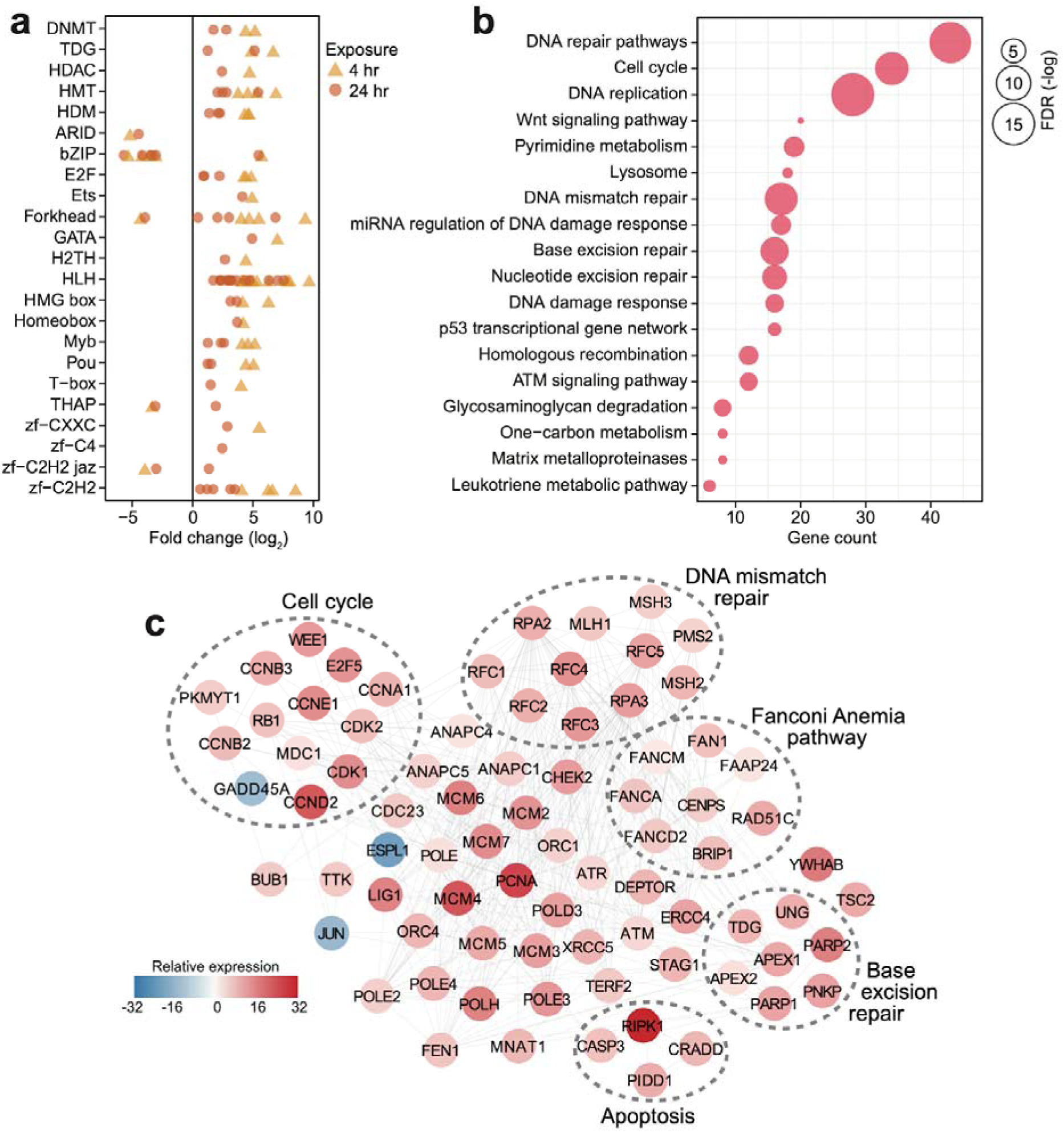
Transcriptional response of *S. caliendrum* to thermal stress. (A) Differentially expressed members of gene regulatory families at 4 hr (yellow triangles) and 24 hr (orange circles). (B) KEGG pathways enriched in the set of differentially expressed genes at 4 hr or 24 hr timepoints. Gene counts (x-axis) and FDR-adjusted *p*-values (bubble size) are shown. (C) DNA damage response network in *S. caliendrum* thermal stress response at 4 hr exposure. Relative expression of genes was computed as the sum of TPM values relative to time-matched control samples. The network is based on human protein–protein interactions.

### Activation of the DNA damage response in *S. caliendrum*

Pathway enrichment analysis of differentially regulated genes in *S. caliendrum* revealed links to cellular responses to DNA damage (Fig. 3B, Table S20-21). Reconstruction of the protein interaction network for genes related to the DNA damage response in *S. caliendrum* revealed upregulation of DNA damage sensors, *ATM* and *ATR* kinases, along with *CHEK2* kinase (Fig. 3C, Table S22), which set off checkpoint-mediated cell cycle arrest, DNA repair activation, and apoptosis *via* the p53 pathway (Blackford and Jackson 2017). Negative regulators of entry into S-phase (*RB1*) and M-phase (*WEE1* and *PKMYT1*) were also upregulated, further indicating cell cycle arrest despite downregulation of the growth arrest and DNA damage-inducible protein (*GADD45A*) and upregulation of cyclin-dependent kinases (*CDK1/2*), cyclin regulatory subunits (*CCND2*, *CCNE1*, *CCNA1*, *CCNB2/3*), and E2F transcription factor (*E2F5*). Cell cycle arrest may facilitate activity of DNA repair mechanisms, as evidenced by upregulation of the mediator of DNA damage checkpoint protein 1 (*MDC1*) (Abraham 2001).

There was also increased expression of genes implicated in DNA damage surveillance and removal, including players in mismatch repair, base excision repair, nucleotide excision repair, homologous recombination, alternative end-joining, and trans-lesion synthesis (Fig. 3C, Table S22) (X. Li and Heyer 2008; Hakem et al. 2012; Bienko et al. 2010; Krokan and Bjørås 2013). This was accompanied by activation of DNA helicases (*MCM2/4/5/6*), DNA polymerase subunits (*POLD3, POLE, POLE2/3/4, POLH*), proliferating cell nuclear antigen (*PCNA*), DNA ligase (*LIG1*), and flap endonuclease (*FEN1*), which ensure high-fidelity DNA re-synthesis and ligation (Timson, Singleton, and Wigley 2000). Activation of the Fanconi Anemia pathway, which coordinates classical DNA repair pathways (Moldovan and D’Andrea 2009), suggests a well-orchestrated deployment of DNA repair mechanisms in *S. caliendrum* under stress.

High levels of DNA damage block mitotic exit through the spindle assembly checkpoint (SAC) (Mikhailov, Cole, and Rieder 2002). The DNA damage network of *S. caliendrum* (Fig. 3C) revealed downregulation of separin (*ESPL1*) and activation of SAC components, including the mitotic checkpoint serine/threonine-protein kinase (*BUB1*) and dual specificity protein kinase (*TTK*), which inhibit the anaphase promoting complex (*ANAPC1/4/5* and *CDC23*) (Overlack, Krenn, and Musacchio 2014). Increased expression of p53-induced death domain-containing protein 1 (*PIDD1*) and an executioner caspase (*CASP3*), along with associated adapter proteins (*CRADD* and *RIPK1*), signals activation of apoptosis, which is likely if DNA lesions remain unrepaired.

### Expression patterns of stress response genes across species

Comparison of SRG transcript abundance revealed higher basal expression in *F. colemani, M. digitata,* and *A. digitifera* relative to the median expression of all transcripts in each species (Fig. 4A). Around 53-64% of SRGs in these corals may therefore be considered frontloaded (1 745 in *F. colemani*, 1 835 in *M. digitata*, and 2 063 in *A. digitifera*). Notably, SRG levels in these species remained stable even under acute thermal stress, with average fold change ranging from 0.21 to 0.58 (Fig. 4B, Table S23-25). These frontloaded genes include heat shock proteins (HSP70) and efflux pumps (ABC transporters and ion transport proteins), as well as SRGs engaged in redox (aldehyde dehydrogenases, cytochrome p450, aldo/keto reductases, peroxidases, thioredoxins, and glutaredoxins) and conjugative (glutathione S-transferases and sulfotransferases) biotransformation (Fig. 4C). Ferritin and apolipoproteins, which have been shown to correlate with enhanced oxidative stress tolerance (Fischer et al. 2013; Granados-Cifuentes et al. 2013), are among the most highly expressed SRGs in *F. colemani* (*FRIS* and *VIT*), *M. digitata* (2 *FRIS*, *APLP*, and *APOB*), and *A. digitifera* (2 *FRIS* and *VIT*) (Table S23-25). Immune receptors (GPCRs, LDL receptors, SRCRs, immunoglobulins, NACHT-, LRR-, Death-, CARD- and TIR-containing proteins) and wound healing genes, such as ECM components (cadherins, collagen, fibronectins, laminins, thrombospondins, and von Willebrand factors), signaling molecules and receptors (activins, galactose binding lectins, C-type lectins, TGF beta, and TNFs), and MH2 transcriptional regulators, also showed stable high expression under the tested conditions (Table S26). It is worth noting that only 27% of SRGs (1 164 genes), including chemical stress response gene families, are frontloaded in *S. caliendrum* (Fig. 4B-C, Table S26-27).

**Fig 4.**
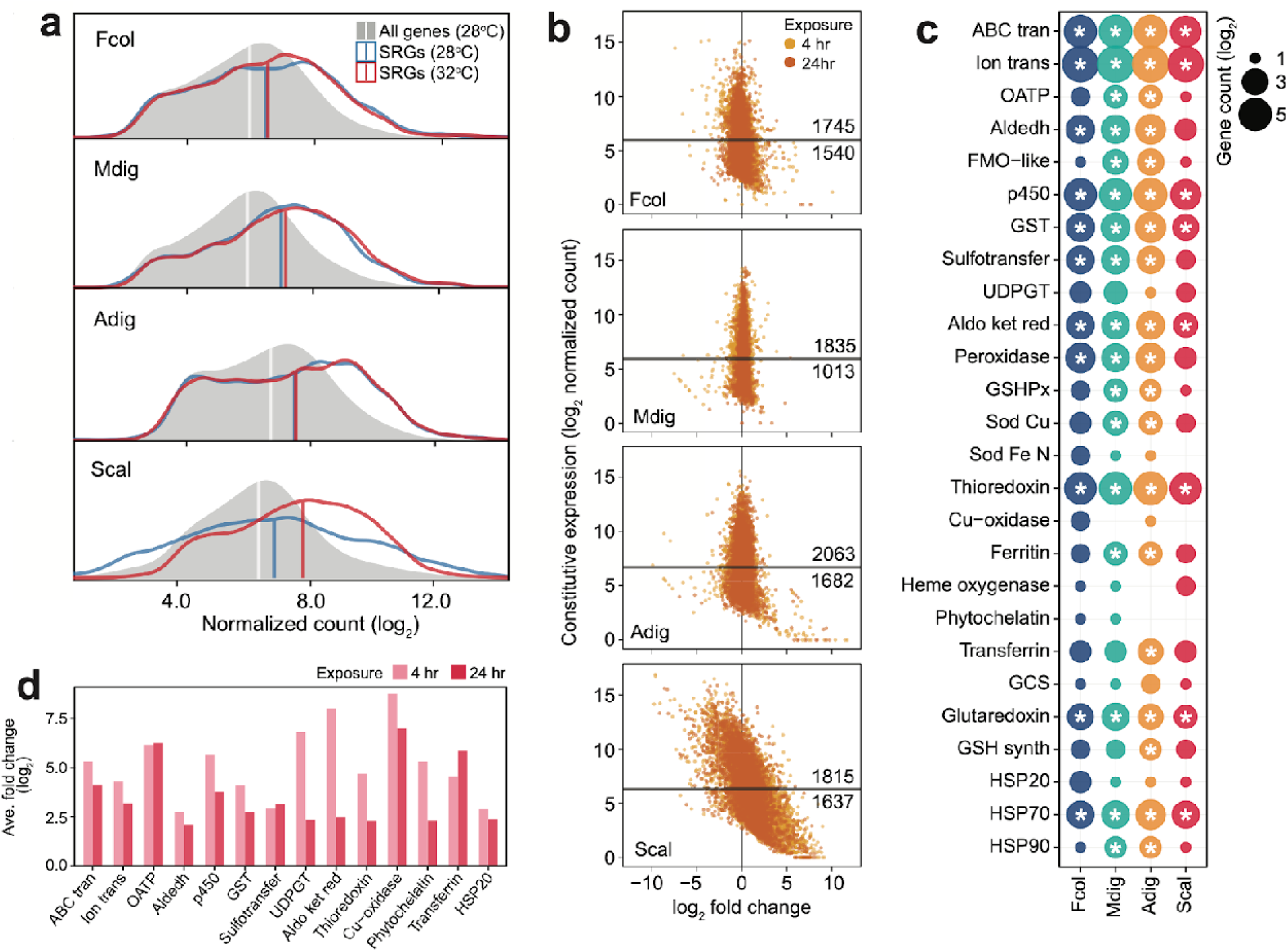
Expression patterns of stress response genes. (A) Expression distribution of all genes at 28 °C (gray shaded area), SRGs at 28 °C (blue line) and 32 °C (red line). Vertical lines indicate median values for each distribution. (B) Expression dynamics of SRGs showing constitutive expression at 28 °C (y-axis) and fold change under thermal stress (x-axis). Y-intercepts indicate median values and numbers above and below this line denote SRGs with higher or lower constitutive expression, respectively, relative to the median. (C) Frontloaded chemical stress response genes. Only genes with normalized counts higher than the transcriptome median expression and with log_2_FC < |2| in the heated treatment were considered frontloaded. Bubble size indicates the number of frontloaded genes and asterisks denote enrichment relative to the total number of peptides in each species. (D) Average fold change of upregulated genes with low basal expression in *S. caliendrum* under thermal stress. Only chemical SRGs are shown.

Unlike in the other corals, SRGs in *S. caliendrum* showed a more dynamic shift in expression under heat stress (ave. |log_2_FC|: 4 hr = 1.70, 24 hr = 1.15) (Fig. 4A-B, Table S27). This was even more apparent for SRGs with basal expression lower than the median for the transcriptome (n = 1 637, 47.42%). These genes showed a greater shift in expression (ave. |log_2_FC|: 4 hr = 2.69, 24 hr = 1.87) (Fig. 4B), with some (n = 524) exhibiting drastic upregulation (log_2_FC ≥ 2) at both 4 hr (ave. log_2_FC = 4.42) and 24 hr (ave. log_2_FC = 3.39) timepoints (Table S28). These lowly expressed but stress-responsive *S. caliendrum* genes are comprised of protein families that are typically frontloaded in *F. colemani*, *M. digitata*, and *A. digitifera* (Fig. 4D, Table S28). Other SRGs in *S. caliendrum* (HSP20, organic anion transporter polypeptides, multicopper oxidases, phytochelatin, transferrin, UDP-glucoronosyl and UDP-glucosyl transferase), cell adhesion (fibroblast growth factors, fibronectins, hemopexin, integrins, and nidogen-like) and innate immunity (inhibitor of apoptosis domain, DEAD, lipoxygenases) proteins also exhibited similar expression patterns (Fig. 4D, Table S28).

## Discussion

In this study, we sought to understand the molecular mechanisms underlying differences in the physiological response of corals to a common thermal stress regime. Using transcriptome sequencing, we identified gene expression signatures of stress responses in these corals. We discovered that gene expression responses, particularly for known stress response genes (SRGs), varied across species, indicating that different corals have distinct strategies to combat thermal stress.

### Variation in SRG copies among corals

Previous comprehensive analysis of genomic and transcriptomic data of diverse corals (Bhattacharya et al. 2016; Zhang et al. 2019), along with the four species from our study, reveals the presence of common stress-related pathways. Notably, different corals possess varying numbers of stress-related genes, suggesting that certain species are equipped with a more diverse set of SRGs (Cunning et al. 2018; Shumaker et al. 2019; Ying et al. 2018; Voolstra et al. 2017; Robbins et al. 2019). Gene family expansion often gives organisms an adaptive advantage as the presence of multiple gene copies may allow for functional diversification or for generation of more gene products (Zhang et al. 2019; Dougan et al. 2024). Expansion of gene families involved in cellular signaling, stress response pathways, and immunity has been reported in the genomes of certain species, including *Pocillopora acuta*, *Stylophora pistillata*, and *A. digitifera* (Cunning et al. 2018; Voolstra et al. 2017). The presence of multiple copies of innate immunity genes could support greater specificity of microalgal endosymbiont recognition (Emery, Dimos, and Mydlarz 2021; Noel et al. 2023) and the ability to recognize and mount responses against pathogens (Shinzato et al. 2011; Baumgarten et al. 2015; Gittins et al. 2015; Alderdice et al. 2022). Higher copy numbers of heat shock proteins (Ying et al. 2018) and fluorescent proteins (Dizon et al. 2021) in massive corals, such as *F. colemani*, could help them better respond to environmental stressors, thereby increasing their chances of survival and reproduction during bleaching events or under thermal and acidification stress (Marshall and Baird 2000; Da-Anoy, Cabaitan, and Conaco 2019; Tañedo et al. 2021).

### Transcriptome plasticity and frontloading of SRGs as an adaptive strategy

The capability of the coral host to modify gene expression in response to environmental stress is critical for recovery and survival (Franssen et al. 2011; Seneca and Palumbi 2015). Global transcriptome change or transcriptome plasticity is a mechanism that promotes the activation of pathways that mediate protection of cellular components or repair of cellular damage (Kenkel and Matz 2016; Studivan, Milstein, and Voss 2019; Karl D. Castillo et al. 2024; Drury et al. 2022; Armstrong et al. 2023). Corals that thrive in highly variable environments typically exhibit greater transcriptome plasticity compared to corals from more stable environments (Kenkel and Matz 2016). One of the most notable observations in our study was that the corals showed different responses to acute thermal stress exposure. While *F. colemani, M. digitata*, and *A. digitifera* exhibited no visible bleaching and little change in gene expression, *S. caliendrum* showed a rapid shift in gene expression profile prior to the onset of bleaching (Da-Anoy, Cabaitan, and Conaco 2019). This change in expression after just 4 hr of exposure reflects rapid activation of the stress response toolkit in *S. caliendrum*. However, about 24% of these differentially expressed host genes remained differentially regulated relative to controls at 24 hr of exposure, indicating a low level of gene recovery. At this point, the coral may have nearly exceeded its thermal response limit and exhausted its cellular resources, which could explain the bleaching observed upon further exposure (Da-Anoy, Cabaitan, and Conaco 2019). A similar response has been reported in the corals, *S. pistillata* (Savary et al. 2021) and *A. hyacinthus* (Thomas et al. 2019), where failure to return to baseline levels of gene expression resulted in low survival likely due to depletion of energy resources.

Notably, the susceptibility of *S. caliendrum* to thermal stress contrasts with the broader understanding that transcriptome plasticity often enhances coral resilience (Kenkel and Matz 2016). Despite rapid gene expression changes indicating a high degree of plasticity, these responses were insufficient to prevent bleaching. This suggests that while *S. caliendrum* can activate protective gene pathways, the rapid onset of these changes may also signal an impending threshold of physiological tolerance. The inability to recover baseline levels of gene expression could deplete cellular resources, which may explain the susceptibility of this species to thermal stress. This finding emphasizes that transcriptome plasticity, while generally advantageous, has its limitations, particularly in species facing extreme or prolonged environmental stressors (Savary et al. 2021; Thomas et al. 2019; Barshis et al. 2014). Across all species examined, the transcriptional response to thermal stress was more pronounced in the coral host than in the algal symbionts. This aligns with previous reports suggesting that the host may insulate its symbionts from external stressors (Barshis et al. 2014; Davies et al. 2018; Kaniewska et al. 2015). However, we expect that more gradual or longer durations of exposure may elicit different responses than what we observed in the acute thermal stress regime used in this study.

In contrast to *S. caliendrum*, *F. colemani*, *M. digitata*, and *A. digitifera* exhibited greater tolerance to acute thermal stress. These species showed higher baseline expression of SRGs at ambient temperature and the expression of these genes remained stable even at elevated temperature. This provides evidence for transcript frontloading, an adaptive molecular response where protective genes are expressed by the cell at higher levels in anticipation of possible stress exposure (Barshis et al. 2013). For example, corals living in warmer or more variable conditions constitutively upregulate a set of genes that are usually only expressed during heat stress (Barshis et al. 2013; Fifer et al. 2021). Frontloading of SRGs is a likely adaptation for corals in habitats that frequently experience stressful conditions and could support tolerance to thermal fluctuations (Bay and Palumbi 2017; K. D. Castillo and Helmuth 2005; Kenkel and Matz 2016; Mayfield, Fan, and Chen 2013; Seneca and Palumbi 2015).

### DNA damage repair in thermally sensitive corals

Functional analysis of differentially expressed genes in *S. caliendrum* revealed evidence for upregulation of protein degradation, transport, DNA damage repair, homeostasis, detoxification, and catabolic processes, which are associated with mechanisms involved in maintaining stable conditions within the coral holobiont. This emphasizes the importance of removing damaged molecules and maintenance of protein conformation and activity (Kenkel et al. 2011; Leggat et al. 2011; Meyer, Aglyamova, and Matz 2011b; Rodriguez-Lanetty, Harii, and Hoegh-Guldberg 2009; Rosic et al. 2011) to properly regulate cellular processes that would then allow the recovery and survival of the organism (Traylor-Knowles et al. 2017).

Maintenance of DNA integrity is essential for homeostasis and survival both in stable and stressful environments (Giglia-Mari, Zotter, and Vermeulen 2011). DNA damage is a consequence of high oxidative stress through the generation of reactive oxygen species (ROS) that results in DNA, protein, and lipid damage (Lesser and Farrell 2004; Lesser 2005) and dysbiosis between coral host and algae (Inoue and Kawanishi 1995; Keyer, Gort, and Imlay 1995; Keyer and Imlay 1996). We found that the thermally sensitive coral, *S. caliendrum*, exhibited higher expression of DNA damage repair genes in response to stress, which indicates that the coral is capable of mobilizing cellular mechanisms to regain homeostasis and could possibly survive brief periods of acute stress. Whether these same cellular mechanisms would support coral survival under gradual or periodic thermal stress events remains to be determined.

## Conclusion

This study contributes valuable genetic information on four common coral species in the Indo-Pacific. We show that these corals possess all the typical genes required to mount an appropriate stress response and are able to express these, especially during stress conditions. Notably, the corals showed very different responses to a common thermal regime. Species that were resistant to thermal stress showed signs of readiness in terms of gene complement, with expanded SRG families, and also in terms of frontloading, with SRGs constitutively expressed in anticipation of stress. The combination of possessing these genetic toolkits and being able to regulate expression in response to various stress events can spell the difference between survival and bleaching. This variation in adaptive strategies could reflect differences in abundance and distribution of these corals on reefs that experience different thermal fluctuations. Future studies to evaluate the potential of these genes and expression patterns as biomarkers for coral thermotolerance will help us better understand the impact of thermal stress across different locations.

## Acknowledgments

We thank the Bolinao Marine Laboratory staff, Marcos Ponce, Ronald De Guzman, Romer Albino, Michael Nada, Jake Baquiran, and Mikhaela Tañedo, for field support and assistance with the thermal stress setup. This study was funded by the Department of Science and Technology Philippine Council for Agriculture, Aquatic and Natural Resources Research and Development (DOST-PCAARRD) Coral Genomics Program to CC.

## Data availability

*De novo* transcriptome assemblies used in this study are deposited at DDBJ/EMBL/GenBank under the following accessions: GIVN00000000 (*F. colemani*), GIVM00000000 (*M. digitata*), GIVI00000000 (*A. digitifera*), and GIVG00000000 (*S. caliendrum*). Sequence reads are available in NCBI under the following BioProject accessions: PRJNA422022 (*F. colemani*), PRJNA422015 (*M. digitata*), PRJNA421253 (*A. digitifera*), and PRJNA422012 (*S. caliendrum*). The final non-redundant reference transcriptomes, as well as datasets generated and analyzed in this study are available on Figshare (https://figshare.com/projects/Coral_transcriptomes/185029).

## Author contributions

JPD & CC conceptualized the study. JPD conducted the experiment. JPD, NP, & CC analyzed the data, drafted and finalized the manuscript.

## Supplementary Figure

**Fig. S1.**
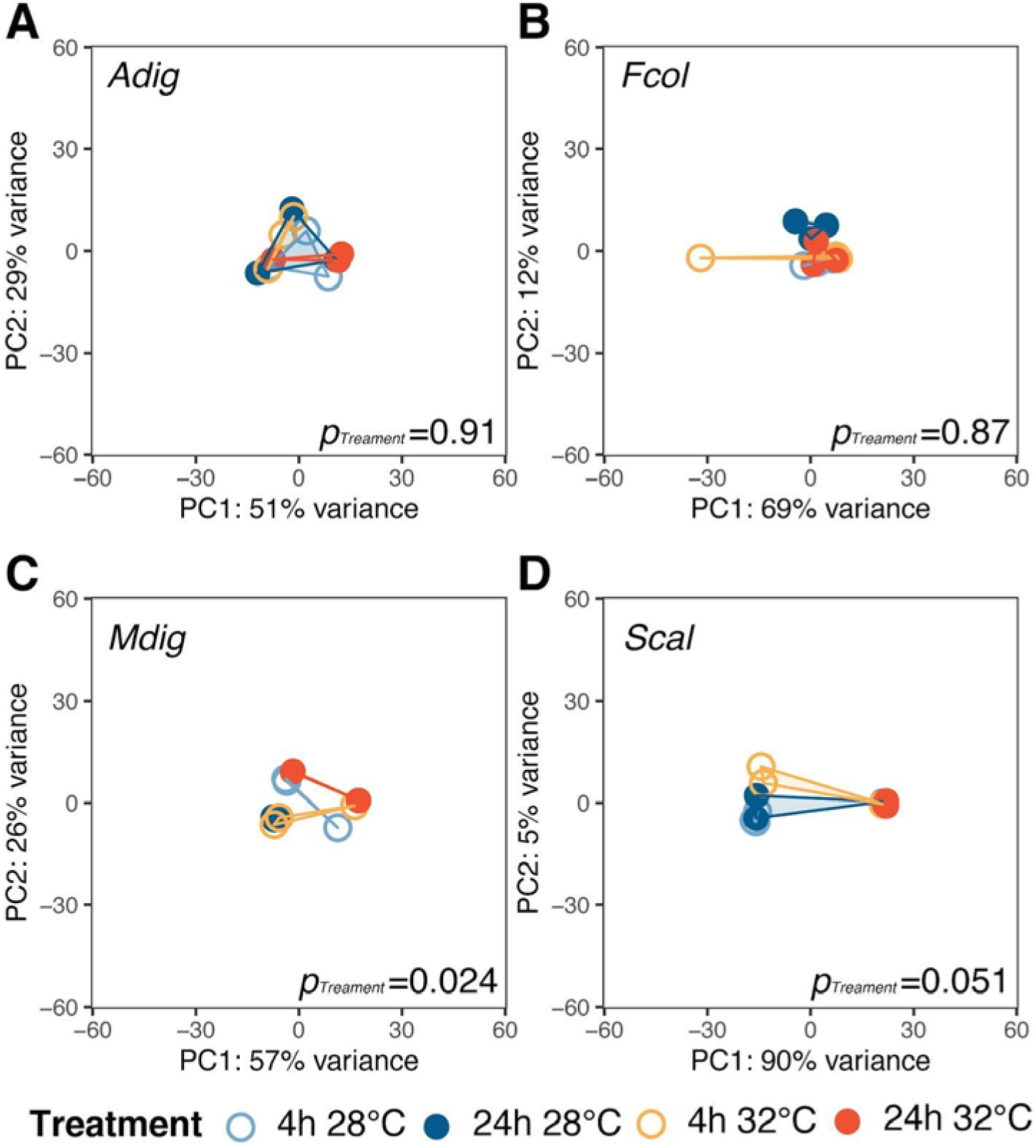
Symbiont transcriptome dynamics under thermal stress. Principal component analysis (PCA) and gene expression plasticity plots of symbiont-derived transcriptome profiles for (A) *F. colemani*, (B) *M. digitata*, (C) *A. digitifera*, and (D) *S. caliendrum* in different treatments. The x- and y-axes represent the % variance explained by the first two principal components.

